# Cardiac oxidative stress monitoring enabled by hierarchical mechanical adaptation

**DOI:** 10.64898/2026.04.15.718718

**Authors:** Bowen Yang, Jianwu Wang, Dong Wu, Zan Chen, Yimei Du, Xiaoxuan Gong, Hong Liu, Yuanyuan Xie, Xiaojun He, Guoliang Hao, Gongxin Wang, Zun Zhang, Kewei Xie, Yong-Xin Wu, Can Cao, Nuan Chen, Pingqiang Cai, Lanxin Xiao, Lei Xie, Haochen Zou, Qunli Lei, Xiao Zhao, Ting Li, Jie Chao, Zhi Jiang, Benhui Hu, Ting Wang, Xiaodong Chen, Lianhui Wang

## Abstract

Soft bioelectronics have advanced cardiac monitoring through electrophysiological tracking, yet this alone cannot resolve the metabolic pathology essential to surgical decision-making. However, real-time molecular sensing on beating hearts remains unresolved due to deformation-induced sensor failure and stress-induced metabolite artifacts. This challenge is exemplified by ischemia-reperfusion injury (IRI), a major cardiac surgery complication characterized by reactive oxidative species (ROS) bursts, where true pathological ROS signals being confounded by mechanotransduction-induced ROS artifacts. Herein, we propose an enzymatic cardiac oxidative stress biosensor (E-cardiac) with hierarchical mechanical adaptation: macro-scale biofluid-mediated contact, micro-scale fiber reorganization, and nano-scale enzymatic confinement within gold nanoarches dissipate interfacial stress. This produces ultrathin (∼460 nm), soft (0.79 kPa) E-cardiac with robust electrochemical stability (100% strain), low detection limit (380 nM), rapid adhesion (<3 s), stable biosensing on beating heart, as well as minimal invasive deployment capability. Mechanical analysis and cellular studies confirm mitigated stress-induced ROS and absent PIEZO channel activation. Validated across cardiomyocytes, *ex vivo* tissues, multi-species ischemia models (mouse, rat, rabbit, pig), rat ischemia-reperfusion injury, and Langendorff hearts simulating graded perfusion deficits, E-cardiac quantitatively differentiates IRI severity (sham < ischemia < reperfusion) as well as detecting the “ECG blind window”. The E-cardiac platform provides real-time metabolic feedback for surgical guidance during cardiac procedures, enabling timely intervention before irreversible damage.

## Introduction

In recent decades, soft implantable electronics have transformed cardiovascular disease management by enabling early diagnosis and timely therapeutic interventions^1,2^. While cardiac electrophysiological monitoring has achieved remarkable sophistication^3^, a critical unmet clinical need persists: real-time molecular monitoring^4,5^ during cardiac surgery to guide intraoperative decision-making^6, 7^. Ischemia-reperfusion injury (IRI) exemplifies this challenge, arising inherently when cardiac surgeries require coronary blood flow interruption and restoration.^8^ The reperfusion process triggers massive reactive oxygen species (ROS) generation within minutes, causing oxidative stress, cellular damage, myocardial dysfunction, and increased post-operative complications^9, 10^. Despite these impacts, surgeons lack real-time monitoring tools: ECG detects only late-stage electrical dysfunction while missing early metabolic injury^8^, and is completely absent during cardioplegic arrest^11^; biochemical markers (e.g., cTnI) require *ex vivo* analysis^12, 13^, creating a diagnostic blind period that precludes verification of myocardial perfusion adequacy. Real-time intraoperative ROS monitoring could enable immediate reperfusion injury assessment and guide targeted interventions to minimize oxidative damage.

However, acquiring reliable biochemical signals from dynamic tissues is fundamentally limited by mechanotransduction^14, 15^, as the heart’s dynamic environment imposes compressive stresses for stable adhesion to its wet, slippery cardiac surface^16, 17^, along with cyclic stretching from continuous contractions^18, 19^, which remains unattainable for current bioelectronics landscape (**Supplementary Fig. 1**). While electrophysiological sensors have demonstrated robust operation across a broad spectrum of tissue biomechanics, ranging from quasi-static neural (∼1% strain) and epidermal (30% strain) interfaces to the highly dynamic myocardium (1–2.5 Hz, 30% strain), the resulting mechanical stimuli can modulate tissue biochemistry^20^, inducing endogenous molecular production that interferes with the detection of target biomarkers. Taking the mechanical stress-induced ROS artifact as an example, cardiac myocytes can transduce external mechanical forces^21^ into intracellular biochemical responses via stretch-activated Ca^2+^ mechanochemical signaling pathways^22^, leading to ROS production that interferes with the pathological oxidative stress signals^23^. Although various strategies have been proposed to mitigate interfacial bioelectronic-tissue stress^24, 25, 26^ like using intrinsically stretchable materials^27, 28^ or triggered adhesion strategies^29,30^ and ultrathin films^31, 32^, this challenge becomes even more pronounced in multilayer biosensing systems^33, 34^. Biosensors typically integrate biotransducers, conductive electrode arrays, and substrates with heterogeneous mechanical properties^35, 36^, leading to multiscale stress concentrations^37^ at both interlayer and tissue–sensor interfaces that surpass the capabilities of conventional single-interface solutions^38, 39^.

Here we introduce an enzymatic cardiac oxidative stress sensor (E-cardiac) enabling real-time molecular monitoring on the beating heart during cardiac surgery (**Figure 1a-c**). Applicable in both open-heart and minimally invasive procedures, E-cardiac detects metabolic deterioration in the “ECG blind window” where electrical function remains normal, enabling intervention before irreversible damage (**Figure 1b and c**). The biosensing mechanism is that E-cardiac comprises cross-aligned microarched gold fibers with encapsulated Prussian blue (PB) nanocatalysts converting H_2_O_2_ (selected for its stability and membrane permeability among ROS species^40^) to electrochemical signals. To address the stress-induced mechanotransduction artifact existed in conventional substrate based biochemical sensors (**Figure 1d, e**), we propose a hierarchical mechanical adaptation strategy: macro-adaptive adhesion dissipates contact stress, micro-fiber reorganization redistributes mechanical loads while maintaining electron transfer pathways, and nano-confined catalyst preserves biotransducer-electrode interfacial stability (**Figure 1f**). This design enables interfacial stress <5 kPa, suppressed PIEZO activation (transcriptomic analysis, calcium imaging, and computational simulations) and robust electrochemical stability, ensuring artifact-free H_2_O_2_ quantification during dynamic cardiac monitoring. We validated E-cardiac across biological scales—cardiomyocytes, *ex vivo* tissues, *in vivo* models (mouse, rat, rabbit, pig). E-cardiac successfully distinguishes sham (∼8.5 μM), ischemia (∼50.2 μM), and reperfusion (∼104.1 μM) phases with excellent correlation to gold-standard colorimetric assay. In Langendorff-perfused rat hearts with simultaneous ECG monitoring, E-cardiac detected escalating oxidative stress at 100-200 μM H_2_O_2_ where ECG remained normal, while arrhythmia appeared only at ≥300 μM, demonstrating early detection capability. An 8-channel array enabled spatial mapping of oxidative stress on beating hearts, localizing ischemic regions corresponding to specific coronary territories. Moreover, we further validated minimally invasive via endoscope-guided deployment, scalability (60 channels), and multimodal sensing capability (glucose, lactate, pH), providing comprehensive metabolic assessment for real-time surgical guidance.

**Fig. 1.**
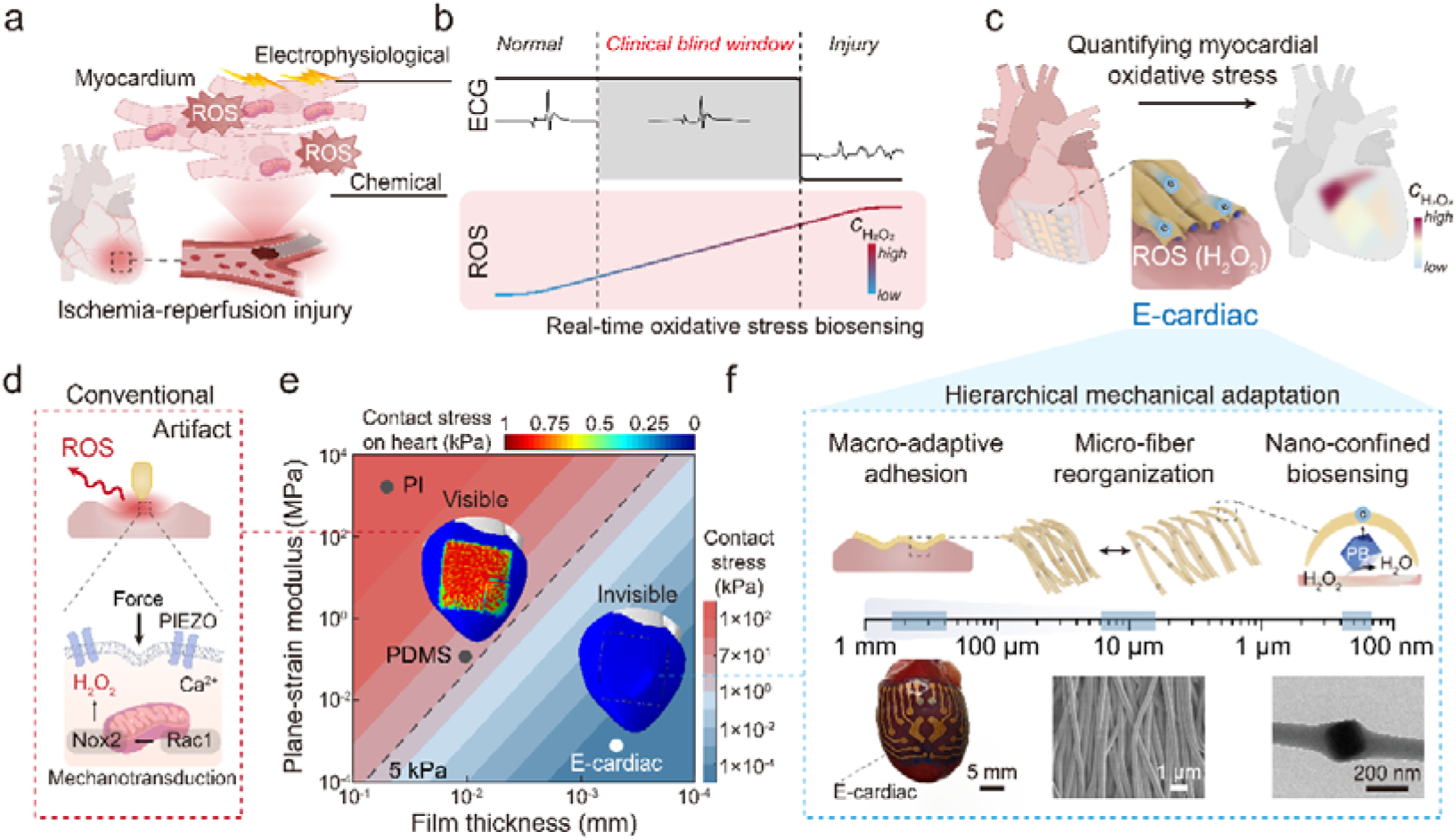
| Hierarchical mechanical adaptation of E-cardiac for high-fidelity oxidative stress monitoring on beating hearts during cardiac surgery. **a**, Schematic of ischemia-reperfusion injury (IRI) where temporary coronary occlusion followed by blood flow restoration triggers acute mitochondrial ROS bursts in the myocardium, with electrophysiological and chemical signals serving as complementary readouts. **b**, Concurrent ECG (black) and real-time HLOL (gradient) monitoring reveal that oxidative stress escalates progressively during the “clinical blind window” where ECG remains normal, providing early metabolic warning that enables intervention before irreversible myocardial damage occurs. **c**, Our E-cardiac enables quantitative myocardial oxidative stress monitoring through conformal epicardial deployment, profiling ROS spatiotemporal distribution during cardiac cycles. **d**, Challenge: mechanical stress can activate PIEZO-mediated mechanotransduction, generating artifactual ROS that confounds true pathological signals through the activation of PIEZO channels involving Ca^2+^ influx and subsequent ROS production. **e**, Contour plot of the simulated contact stress at the electrode-tissue interface for devices with varying plain-strain modulus and thickness, positioning E-cardiac in the ultra-soft (∼0.79 kPa) and ultrathin (460 nm) regime that minimizes contact stress below the 5 kPa threshold, contrasting with conventional flexible electronics substrate materials like PDMS (10 μm-thick) and PI (50 μm-thick). Inset: Finite element simulation showing contact stress distribution for E-cardiac (left, blue region indicating minimal stress) and PDMS control (right, red region indicating high stress concentration) when interfacing with heart tissue. **f**, Hierarchical stress deconcentration mechanism in E-cardiac operating across three scales: millimeter-scale adaptive adhesion, micrometer-scale fiber reorganization, and nanoscale confinement. Digital images show E-cardiac achieving conformal contact with a rat heart (5 mm scale), cross-aligned gold fiber networks (1 μm scale), and nanoconfined PB particles (200 nm scale).

### Design, fabrication and characterization of E-cardiac

Different from conventional layer-by-layer interfacial deposition of biotransducers, we propose a nanoarching strategy where biocatalysts are encapsulated within nano-arched microfiber structures. The key design principle is that mechanical strain is progressively deconcentrated across three hierarchical scales: (upper panel in **Figure 1f**): macro-scale adaptive adhesion through biofluid-mediated conformable contact dissipates contact stress, micrometer-scale fiber reorganization redistributes cardiac beating-induced stress while maintaining stable electron transfer pathways; and nano-catalysts confinement protects enzymatic catalysts within gold microarches from mechanical deformation. The microscopy images show E-cardiac achieving fully conformal contact with wet tissue surfaces (10 mm scale), cross-aligned gold fiber networks (1 μm scale), and nanoconfined PB particles (200 nm scale) (lower panel in **Figure 1f**), corresponding to the aforementioned hierarchical stress deconcentration mechanism.

It is reported that mechanical stress-stimulated ROS production in cardiomyocytes is primarily attributed to activation of mechanosensitive PIEZO channels (**Figure 1d**). This pathway becomes markedly active when interfacial stress exceeds several kPa (8 kPa^41^), triggering Ca^2+^ influx and promoting ROS production. Therefore, mechanically invisible biosensors that minimize interfacial stress are essential for detecting true IRI-related ROS levels without artifacts.

To theoretically predict the contact behavior at the tissue-device interface, we established an analytical model based on an elastic substrate with wavy surface profile^42^, mimicking the microscale tissue topography. We analyzed the contact stress generated by materials with different plane-strain moduli and thicknesses, details given in the **Supplementary Note 1**, **Equation (S7)**. During the adaptation process, E-cardiac induced contact stress well below kilopascal level, whereas PI (50 μm) generated contact stress exceeding this limit and PDMS (10 μm) is almost in the region (**Figure 1e**). Finite Element (FE) analysis further confirms the markedly reduced stress distribution produced by E-cardiac upon contact with the heart surface, in contrast to the elevated pressure (∼1 kPa) observed with a PDMS membrane (the inset in **Figure 1e**, **Supplementary Note 2**). A similar trend was observed in calculations of substrate stretching with maximal 20% strain, where E-cardiac accommodated deformation with minimal contact stress, while PI, and PDMS again exceeded the critical stress threshold (**Supplementary Fig. 2**). These results theoretically illustrate the E-cardiac’s low-stress interfacial behavior due to its softness and ultrathin features, which are essential for eliminating mechanotransduction-induced artifacts for high-fidelity ideal biosensing interface.

To fabricate the E-cardiac, we select PB as a model enzymatic catalyst for H_2_O_2_ sensing (details given in **Supplementary Note 3**), and polyvinyl alcohol (PVA) as the sacrificial core under the microarched gold shell for oxidative stress monitoring under dynamic cardiac conditions. The E-cardiac is fabricated through radial electrospinning of PVA solution containing PB nanoparticles, followed by thermal deposition of gold (**Supplementary Fig. 3 a, b**). The sensing area of working electrodes for the PVA/PB-Au electrodes was defined with an Al_2_O_3_ insulator layer (**Supplementary Fig. 3c-e**). We also developed a patterned 2 × 4 sensor array, enabling continuous spatial mapping of oxidative stress, which allows precise identification of ischemic regions during disease on-set. Digital images show that the obtained sensor array is ultra-lightweight, able to rest on a dandelion (**Supplementary Fig. 4**). As demonstrated in dynamic cardiac environments, E-cardiac can conform to beating rat (**Supplementary Video 1**) and pig hearts (**Supplementary Video 2**) within 3 seconds and maintain conformal contact under dynamic heart beating, preserving seamless integration throughout cardiac cycles. This conformability of E-cardiac is expected to result from the hierarchical mechanical adaptation strategy that enables ideal stress-resilient biosensing interfaces.

The optimal PVA concentration (10 wt%) yielded uniform fibers (**Supplementary Fig. 5**) to fabricate the cross-aligned PVA nanofibers with diameters around 180 nm (**Supplementary Fig. 6**). Gold coating further increased fiber diameters to 260 nm ± 20 nm (**Supplementary Fig. 7a**), with energy-dispersive X-ray spectroscopy (EDS) mapping showing overlapped Fe (from PB) and Au signals along the fiber direction, validating the formation of PB-encapsulated gold tubular structures (**Supplementary Fig. 7b**).

### Macro-scale mechanical adaptation to deconcentrate adhesion stress

We investigate the stress-deconcentrating adhesion process of E-cardiac to wet tissues through structural and mechanical characterization. To visualize the conformability transition, we incorporated fluorescent cyanine 5 in PVA fibers and monitored their behavior on the surface wrinkles of tissue-mimicking templates using a confocal laser scanning microscope (CLSM). The conformability of the E-cardiac film was validated using fluorescence and bright-field imaging, which exhibited a clear transition from an opaque state to a transparent state, indicating intimate contact with the wrinkled replica (**Figure 2a**). The Z-stack scanning and reconstructed images further revealed structural evolution: the initially flat film in its dry state adapted to surface contours upon wetting, precisely matching wrinkle depths around 5 μm (**Supplementary Fig. 8**). E-cardiac demonstrated superior conformability to complex structures, including skin replica microstructures (**Supplementary Fig. 9**) and gel microspheres (diameter ∼300 μm, **Supplementary Fig. 10**), outperforming conventional soft substrates like PDMS film.

**Fig. 2.**
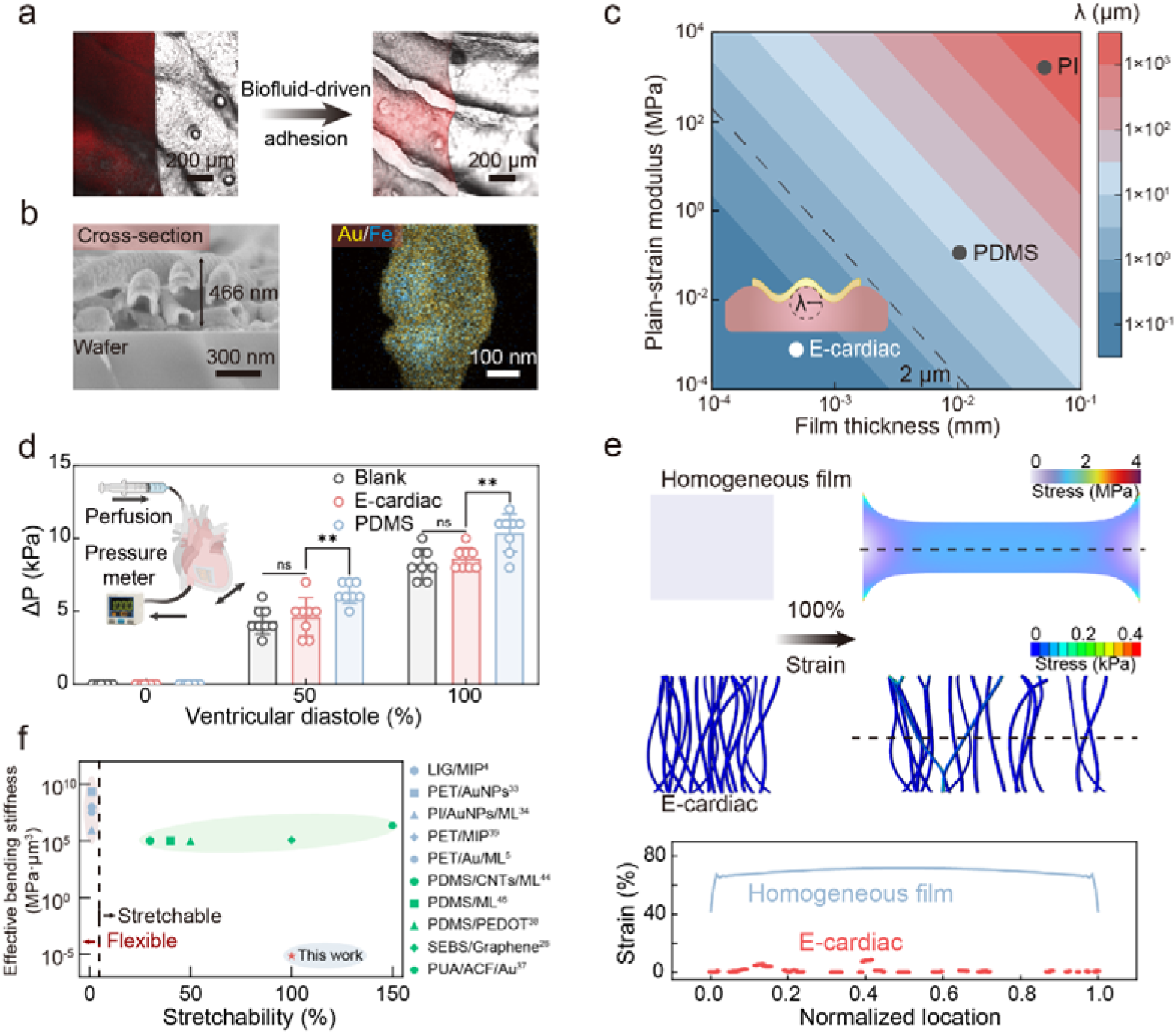
| Morphology and mechanical analysis of stress-deconcentrated E-cardiac during tissue adhesion and dynamic deformation. **a**, CLSM images of PVA fibers at dry state (left) and after water-driven adhesion (right) to skin replica, demonstrating conformal contact to surface topography. **b**, Micromorphology of E-cardiac showing cross-sectional view and element distribution revealing cross-aligned microarched Au fibers with PB nanoparticles encapsulated after biofluid treatment. **c**, Theoretical analysis of stress-deconcentrated conformal contact. Contour maps showing relationships between plane-strain modulus, film thickness, and contact radius (1 kPa pressure). Insets illustrate biofluid-driven adaptive adhesion process on spherical. **d**, Simulated diastolic pressure measurements during cardiac motion simulation (inset: experimental setup with PBS circulation). E-cardiac attached heart shows negligible pressure difference from bare heart, while PDMS attachment significantly increases mechanical constraint. **e**, Finite element simulations comparing strain distributions in substrate (up) and E-cardiac (down) under 100% strain, demonstrating minimal strain (<0.5%) at gold microarch junctions. **f**, Comparison of effective bending stiffness versus stretchability for the developed E-cardiac (this work) against various reported flexible and stretchable biosensors,^4, 5, 26, 33, 34, 37, 38, 39^ highlighting its advantages in the biosensor regime.

To quantify the mechanical changes during this process, we performed *in situ* Young’s modulus measurements using Piuma nanoindentation probes (details given in **Supplementary Note 4**). The modulus of E-cardiac film significantly decreased from 1.27 MPa to 0.79 kPa after PVA dissolution (**Supplementary Fig. 11**). Assessment of the mechanical impact on tissue surfaces (detailed image shown in **Supplementary Fig. 12a**) revealed that tissues with attached E-cardiac maintained modulus values close to native cardiac tissue (16.0 ± 1.7 kPa), whereas conventional elastomers such as PDMS induced significantly elevated modulus exceeding 1 MPa (**Supplementary Fig. 12b**). E-cardiac attachment to other soft organs, including liver (4.2 ± 0.1 kPa), spleen (25.1 ± 1.9 kPa), lung (4.1 ± 0.2 kPa), and kidney (15.4 ± 1.6 kPa) (**Supplementary Fig. 12c**), similarly maintained modulus values comparable to native tissue without detectable elevation. These validations of biomechanical compatibility across diverse models and organ tissues indicate that the E-cardiac device causes minimal perturbation at soft, dynamically active biological interfaces. Moreover, it is also noted that dissolved PVA provided interfacial stability, generating an adhesion force of approximately 0.015 N to a wet tissue-mimic surface via 90-degree peel testing (**Supplementary Fig. 13**).

The fibrous integrity during PVA dissolution was investigated to confirm preservation of the sensing architecture. Cross-sectional SEM images show that the E-cardiac thickness decreases from ∼2.60 µm (**Supplementary Fig. 14**) to ∼460 nm following the PVA dissolution process, while individual fibers adopt micro-arched architectures (**Figure 2b**, left). SEM imaging further confirms that the cross-aligned fiber network is preserved throughout dissolution **(Supplementary Fig. 15a, b**), a crucial feature for maintaining both mechanical resilience and electrical continuity. After PVA removal, top-view SEM images reveal that initially loosely stacked fibers become more densely packed, and fiber orientation analysis by fast Fourier transform (FFT) showed no significant change in the alignment structure of E-cardiac before and after PVA removal **(Supplementary Fig. 15c, d**). And PB nanoparticles remain securely confined within the fibrous structure, as confirmed by EDS (**Figure 2b**, right). These results collectively demonstrate that PVA dissolution under wet conditions enables E-cardiac to transition from a rigid state to a soft, hydrated phase, during which PB nanoparticles remain confined to the electrode-tissue interfaces.

To elucidate the underlying mechanisms of the adhesion process, we employed an epidermal contact model, characterized by a sinusoidal profile with a 2 μm wavelength and amplitude, comparable to skin wrinkles. This model correlates the work of adhesion with membrane properties and surface geometry (see **Supplementary Note 1**). Based on the aforementioned thickness and modulus, we used the model to determine the critical surface wavelength *λ*_0_ of the heart that allows a thin membrane with varying plane-strain modulus and thickness to conformally adhere. Noted that PVA dissolution creates an *in situ* molecular adhesive layer that provides an effective work of adhesion of 0.49 J/m^2^ (**Supplementary Fig. 13**). This value significantly exceeds the threshold required for electrode-tissue adhesion. Based on this adhesion energy, theoretical predictions indicate that E-cardiac can achieve spontaneous conformal contact with features down to 0.09 μm in wavelength (**Figure 2c**). In contrast, conventional materials such as PDMS (*E* ≈ 0.1 MPa, thickness ≈ 10 μm) and PI (*E* ≈ 1.6 GPa, thickness ≈ 50 μm) are limited to conforming with much larger features (*λ*_0_ > 11 μm and 1.3 mm, respectively). These findings demonstrate that hierarchical mechanical adaptation realizes a significant reduction in both modulus and thickness, combined with *in situ* formation of an adhesive layer, enabling stress-mitigating conformal contact with wet cardiac tissue.

### Microscale mechanical stress dissipation under heart beating

The dynamic beating of the heart imposes continuous mechanical stress that extends beyond the initial adhesion process. The E-cardiac, with its cross-aligned microarched structure, is designed to accommodate this dynamic tissue deformation. We tested the integration of E-cardiac with tissue-mimicking substrates (Young’s modulus: 180 kPa–1.24 MPa) to evaluate its effect on mechanical properties under stretching. E-cardiac was directly attached to mimic tissues through water-driven adhesion, while PDMS films (100 μm) as a control were attached using partially cured PDMS as an adhesive. The nearly identical stress–strain curves of tissue-mimicking substrates before and after E-cardiac adhesion indicate that E-cardiac exerts negligible mechanical impact on the underlying tissue under the beating heart, whereas conventional PDMS attachment significantly increases the effective modulus (**Supplementary Fig. 16**).

We further simulate the dynamic tissues with an isolated heart with circulating liquid to mimic the beating heart, and monitored pressure during this process (the setup shown in the inset of **Figure 2d**). Upon equivalent PBS solution injection, hearts with E-cardiac attachments maintained their native pressure response, whereas PDMS attachments significantly restricted motion, leading to increased internal pressure (**Figure 2d**). Similarly, in an Ecoflex tube model simulating blood vessel expansion (**Supplementary Fig. 17a**), E-cardiac preserved the vessel’s natural pressure dynamics during expansion, while PDMS-induced restriction resulted in elevated pressure (**Supplementary Fig. 17b**). These results demonstrate that E-cardiac’s cross-aligned network structure accommodates tissue deformation and distributes interfacial stress while preserving natural cardiac contractility.

We conducted FE simulations to investigate the strain distribution in conventional soft substrates compared with E-cardiac, as detailed in **Supplementary Note 2**. Under both 50% and 100% applied strain, E-cardiac exhibited negligible stress on individual microarched fibers. The strain energy was primarily dissipated through network reorganization (**Figure 2e**). In contrast, a homogeneous film (e.g., PDMS) showed significant stress accumulation across the film, reaching 1.26 MPa under 50% strain. Correspondingly, the bottom plot shows the strain distribution extracted along the horizontal dashed line across the film thickness. It reveals that homogeneous films underwent highly localized deformation, with strain exceeding 60% in the necking region. In contrast, the E-cardiac maintained minimal strain levels (<10%) across most locations, suggesting its ability to redistribute and buffer mechanical loads via microscale architectural rearrangement. These results highlight the superior mechanical adaptivity of E-cardiac in dynamic environments.

We also compare the mechanical properties of our E-cardiac with reported biosensing electrodes. It is revealed that the effective bending stiffness of E-cardiac is reduced by three orders of magnitude compared to conventional biosensors with layer-by-layer configurations (**Figure 2f**, **Table S1**). This pronounced mechanical compliance aligns with FE simulation predictions, confirming that hierarchical mechanical adaptation through stress-mitigating adhesion and microarched network reorganization enables effective stress dissipation in dynamic cardiac tissues.

### Negligible ROS artifacts at the E-cardiac and tissue interface

Thanks to the inherent stress-deconcentrating properties, E-cardiac is expected to effectively minimize mechanosensitive pathway activation, thereby reducing artifactual ROS generation. To validate, we conducted parallel experiments to examine the cellular ROS response and mechanoresponsive PIEZO1 channel-related calcium levels under three conditions: cells interfaced with the E-cardiac system, untreated cells (blank), and cells subjected to 5 kPa pressure^31, 43^ as a control (**Figure 3a, b**).

**Fig. 3.**
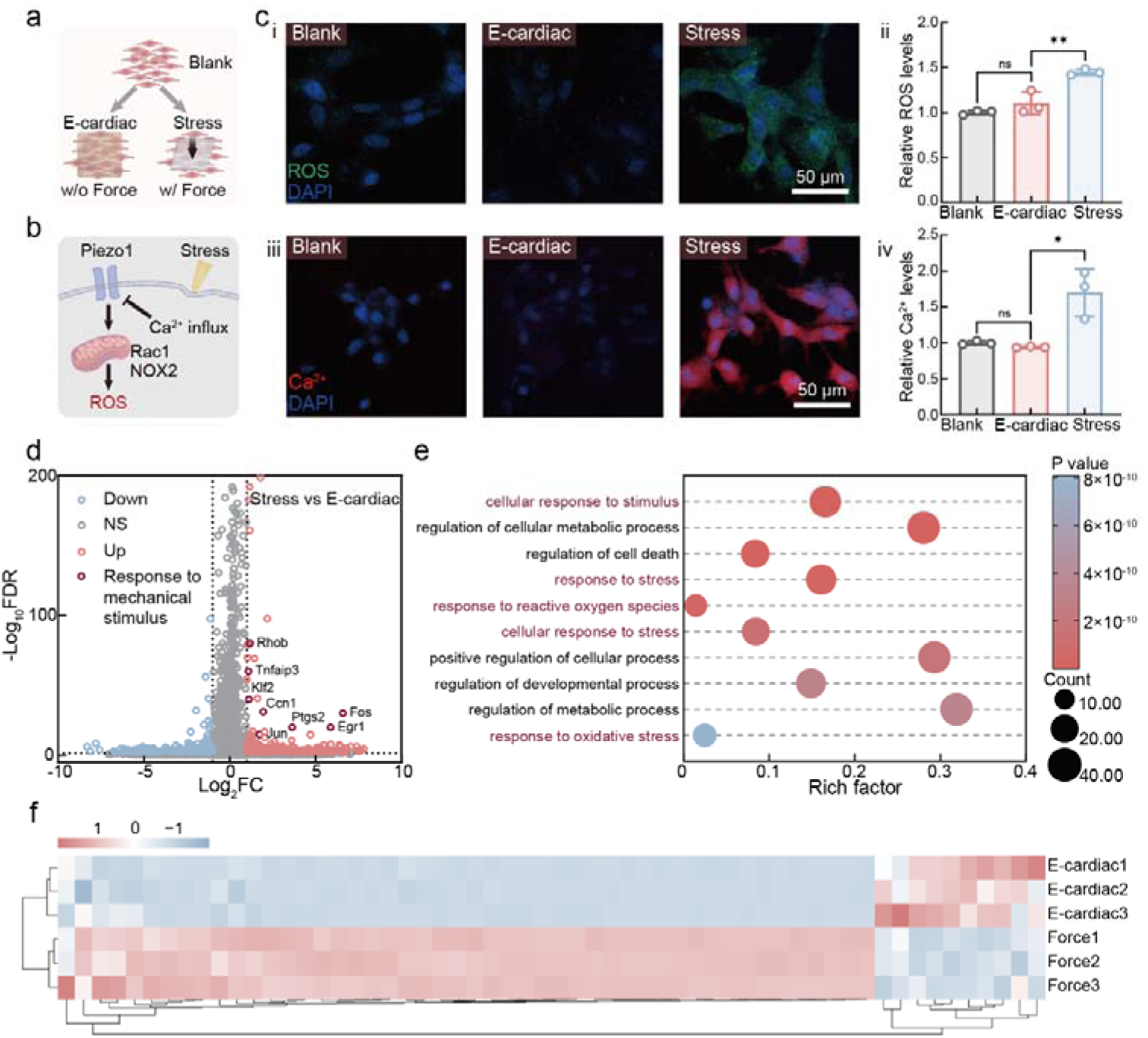
| Cellular mechanisms on the mitigating effect of ROS artifacts at E-cardiac-tissue interfaces. **a**, Schematic comparison of cardiomyocytes responses under three conditions: unstressed cells, E-cardiac interface, and external mechanical stress. **b**, Schematic illustration of mechanosensitive PIEZO channel activation, depicting the cascade from mechanical stress to Ca^2+^ influx and subsequent ROS production. **c**, Representative CLSM images (i, iii) and quantification of fluorescent intensity (ii, iv) comparing unstressed control, E-cardiac interface, and external stress conditions. Intracellular ROS in cardiomyocyte was stained with DCFH-DA (green), calcium ions with Fluo-4 AM (red), and nuclei with DAPI (blue). Error bar for fluorescent intensity in (ii) and (iv) is calculated with three independent groups. **d**, Volcano plot of transcriptomic differences between E-cardiac and Force-treated H9C2 cells, identifying 341 genes significantly upregulated under mechanical stress. **e**, Gene ontology (GO) enrichment analysis of differentially expressed genes (DEGs). DEGs were defined by Benjamini–Hochberg adjusted P < 0.05 and absolute logL fold change ≥ 1. **f**, The GO enrichment heatmap of DEGs between the Stress and E-cardiac groups reveals pronounced transcriptional changes in response to mechanical stress. Genes were clustered by average linkage and Euclidean distance, and values are presented as z-scores normalized per gene.

The cellular response to mechanical stress in both cardiomyocytes (H9C2) and endothelial cells (HUVECs) was investigated with CLSM^44^. We utilized DCFH-DA to visualize ROS levels in cells. Cells interfaced with E-cardiac maintained ROS fluorescence levels comparable to untreated control groups, demonstrating no additional ROS production, while cells exposed to 5 kPa pressure showed significantly elevated fluorescence, as shown by CLSM images and quantitative analysis (**Figure 3c-i, ii**, **Supplementary Fig. 18a**). These results demonstrate that E-cardiac can effectively prevent mechanical stress-induced cellular responses that could interfere with disease-related ROS levels. We further employed Fluo-4 AM as a fluorescent calcium indicator to measure intracellular calcium levels. Cells subjected to mechanical stress exhibited significantly elevated calcium levels. In contrast, E-cardiac-interfaced cells maintained baseline calcium levels comparable to unstressed controls, indicating that E-cardiac effectively mitigated mechanical stress-induced calcium influx, as shown by CLSM images and quantitative analysis (**Figure 3c-iii, iv; Supplementary Fig. 18b**).

Furthermore, we performed flow cytometry analysis to quantify the cell populations based on ROS and calcium levels for both cardiomyocytes and endothelial cells. The distribution of DCFH-DA-stained cells treated with E-cardiac showed ROS levels comparable to the blank group (blank group: 2.41% positive cells in H9C2, 17.4% positive cells in HUVEC; E-cardiac group: 2.67% positive cells in H9C2, 17.0% positive cells in HUVEC), while cells exposed to mechanical stress exhibited a significant increase in ROS production (Force group: 15.5% positive cells in H9C2, 48.8% positive cells in HUVEC; **Supplementary Fig. 19a; Supplementary Fig. 20a**). Similarly, flow cytometry analysis using Fluo-4 AM revealed consistent results: both cell types subjected to mechanical stress displayed significantly elevated calcium levels (Force group: 25.1% positive cells in H9C2, 36.9% positive cells in HUVEC). In contrast, cells treated with E-cardiac maintained comparable calcium levels to unstressed controls (blank control group: 3.53% positive cells in H9C2, 1.76% positive cells in HUVEC; E-cardiac group: 3.12% positive cells in H9C2, 1.71% positive cells in HUVEC; **Supplementary Fig. 19b; Supplementary Fig. 20b**). The consistent responses observed across these distinct cell types support a conserved mechanotransduction pathway, in which PIEZO1-mediated calcium influx leads to downstream ROS production. These findings demonstrate that E-cardiac interfaces effectively minimize artifacts in oxidative stress detection and enable reliable monitoring of disease-related ROS in cardiovascular tissues.

We further investigated the cellular response to mechanical stress in cardiomyocytes (H9C2) using transcriptomic profiling. mRNA sequencing was conducted across three experimental groups: Blank (untreated), E-cardiac-interfaced, and force-treated (5 kPa pressure). Differential expression analysis (DESeq2, FDR-adjusted P < 0.05) identified 58 genes significantly enriched in Gene Ontology (GO) terms upregulated in force-treated cardiomyocyte cells compared to E-cardiac-interfaced counterparts (**Figure 3d**), which primarily involved pathways in cellular response to stimulus, response to ROS, cellular response to stress, regulation of metabolic processes, and oxidative stress (**Figure 3e**). Notably, force treatment loading induced robust activation of hallmark mechanosensitive genes, including *Egr1* (5.9-fold) and *Klf2* (1.1-fold), along with Rho signaling activators *Ptgs2* (3.6-fold) and *Rhob* (1.2-fold) (**Figure 3f**). This transcriptional signature suggests activation of NADPH oxidase (NOX)-mediated ROS generation. In addition, force-treated cells exhibited pronounced pro-inflammatory gene expression, including upregulation of *IL-6* (1.2-fold) and *CXCL1* (1.4-fold) through NF-κB/MAPK pathways, as well as fibrotic markers *Ccn1* (2.0-fold), indicating that sustained mechanical stimulation may promote fibrotic remodeling^45^. In contrast, E-cardiac-interfaced cells displayed minimal transcriptomic alterations relative to blank controls, with only 8 differentially expressed genes (**Supplementary Fig. 21a**), and no significant enrichment in mechanosensitive or oxidative stress-related pathways (**Supplementary Fig. 21b and c**). Collectively, E-cardiac effectively prevents mechanical stress-induced cellular activation, thereby ensuring high-fidelity oxidative stress monitoring.

### Nanoconfined enzymatic sensing under dynamic conditions

To validate the ROS monitoring capability, we fabricated H_2_O_2_ sensors using E-cardiac where the enzymatic nanoparticles preconfined in cross-aligned Au microarched fibers as working/counter electrodes and Ag/AgCl as reference electrodes (**Figure 4a**). The system includes an Al_2_O_3_ insulator layer to define the working electrode area and interfaces with measurement electronics through an ultrathin (6 μm thick) flexible Au/PET connector for signal readout. Noted that this architecture is compatible with alternative peroxidase-mimicking nanocatalysts (e.g., CuO, MnO_2_, and Fe_3_O_4_), enzymes (e.g., glucose oxidase and lactate oxidase), and responsive polymers (e.g., polyaniline), demonstrating the generality of our nanoconfinement strategy as validated in subsequent experiments.

**Fig. 4.**
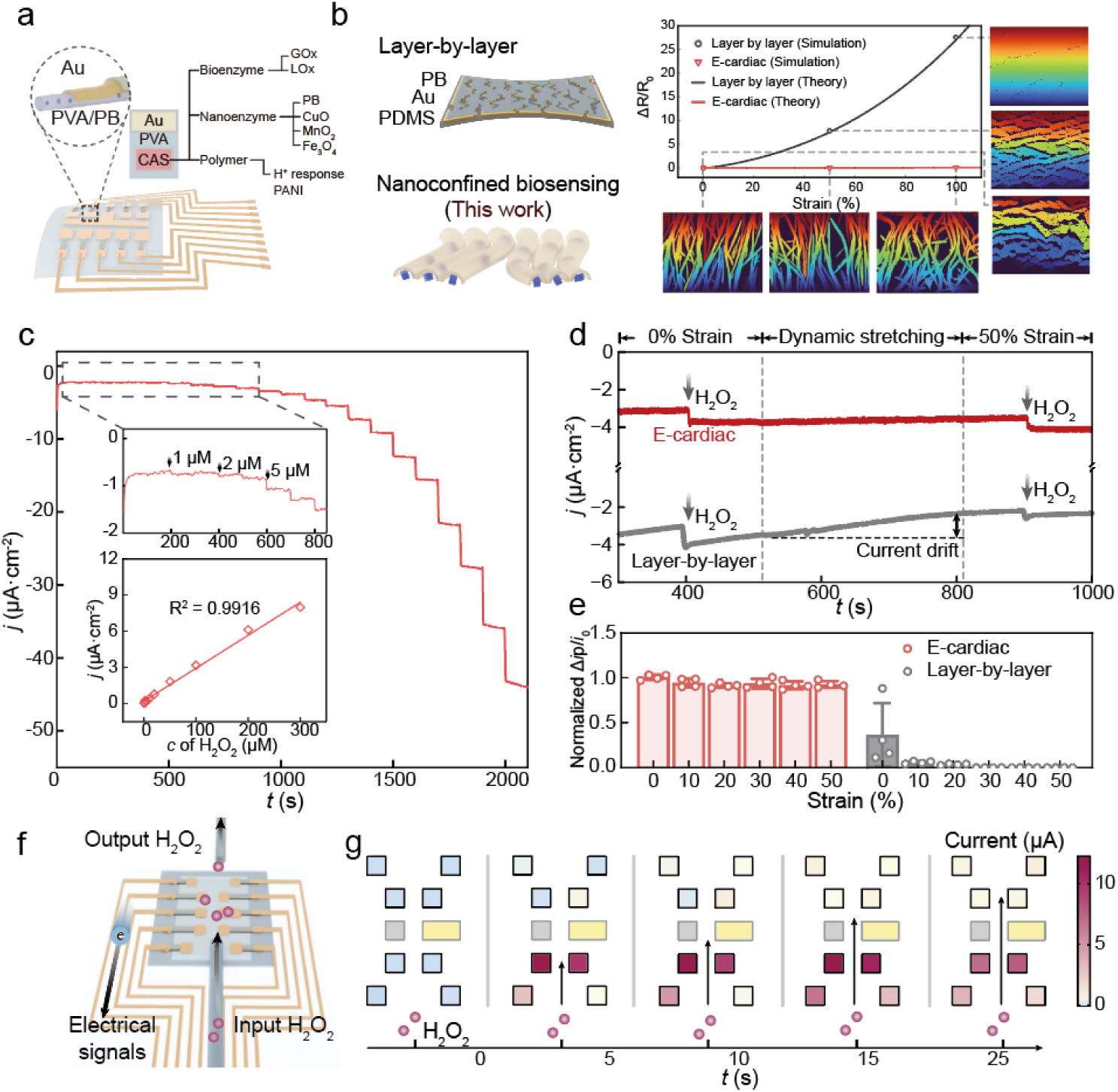
| Robust electrochemical H_2_O_2_ sensing and mapping under stretching conditions. **a**, Schematic of the E-cardiac biosensor array. Inset: a PVA/PB/Au nanofibers where the encapsulated enzymes (applied to mimic enzymes (CuO, MnO_2_, or Fe_3_O_4_), enzymes (GOx and LOx), and H^+^ sensitive polyamine (PANI). **b,** Strain-insensitive electrochemical performance enabled by nanoconfined biosensing architecture. **Left**: Schematic comparing E-cardiac’s nanoconfined PB/Au architecture (bottom) versus conventional layer-by-layer PB/Au/PDMS electrode (top). **Right**: Resistor-network simulations demonstrated that E-cardiac maintains minimal resistance change (Δ*R*/*R*_0_ < 0.08, red line) across 0-100% strain, contrasting with layer-by-layer electrodes (Δ*R*/*R*_0_ > 25, gray line). Corresponding electrical potential distribution maps (right panels) show E-cardiac preserves uniform voltage distribution under strain (0%, 50%, 100%), while layer-by-layer configuration exhibits severe voltage drop and current concentration at defect sites, confirming E-cardiac’s superior electrical stability and reliable H_2_O_2_ sensing during cardiac motion. **c**, Amperometric H_2_O_2_ sensing performance of E-cardiac containing 2.5 wt% PB nanoparticles at −0.05 V vs. Ag/AgCl. The calibration curve demonstrates a strong linear correlation between current response and H_2_O_2_ concentration, indicating the reliability of the E-cardiac. **d**, Amperometric response of E-cardiac (red) and layer-by-layer (PDMS/Au/PB) assembled electrode (gray) to H_2_O_2_ under dynamic strain (0% to 50%). E-cardiac maintained a stable steady-state current in the *i-t* curve under strain, while the layer-by-layer assembled electrode exhibited significant current fluctuations. Schematic illustrating the strain-insensitive electrochemical performance of the E-cardiac (left) and conventional layer-by-layer electrodes (right): The arch structure ensures stable PB confinement and electron transfer even under mechanical deformation. SEM images show the separation of aligned arches under strain, in contrast to the PDMS/Au electrode, which easily cracks under deformation. **e**, Normalized current response to 50 μM H_2_O_2_ under 0-50% incremental strain for E-cardiac (red) and layer-by-layer-structured PDMS/Au/PB electrode (gray). **f, g,** Schematic and spatiotemporal response of 2×4 E-cardiac array with eight working electrodes (colored squares), counter electrode (central gray), and reference electrode (yellow). H_2_O_2_ flow (50 μM, 500 μL/s, arrow indicating flow direction) generates progressive sensor activation (dark purple ∼10 μA), visualizing concentration propagation and demonstrating spatial mapping capability.

Biosensing stability under mechanical deformation is critical for reliable molecular monitoring in dynamic tissues. The nanoconfinement effect in E-cardiac protects catalysts from strain-induced deactivation by preventing PB nanoparticle detachment from nanoarched microconductors (**Figure 4b**, left schematic). Resistor-network simulations further elucidate the underlying mechanism for the improved sensing performance under dynamic conditions. Conventional layer-by-layer Au-PB films exhibit microcracking and severe conductive pathway discontinuity under strain (Δ*R*/*R*_0_ > 25-fold at 100% strain, **Figure 4b** right), whereas E-cardiac’s cross-aligned microfiber architecture maintains intact electrical pathways through dynamic fiber reorganization via sliding, rotating, and reconfiguring rather than fracturing (Δ*R*/*R*_0_ < 0.08-fold). Electrical potential distribution maps confirm that E-cardiac preserves uniform voltage distribution under 0-100% strain, while layer-by-layer electrodes show voltage concentration at defect sites. SEM images demonstrate this controlled deformation mechanism, showing fiber separation while preserving structural integrity at strains up to 100% (**Supplementary Fig. 22**). This architecture maintains highly conductive pathways (502 ± 16 S/cm) along microarched fibers even under 100% strain (**Supplementary Fig. 23**), ensuring stable biosensing performance during continuous cardiac motion. In contrast, conventional layer-by-layer PDMS/Au/PB electrodes exhibited substantial resistance increases under 50% strain, with Δ*R*/*R*_0_ escalating from 6.39 to 21.8 as PB deposition cycles increased from 5 to 10, and complete electrical failure occurring at 15 cycles (Supplementary Fig. 24). Similarly, Ag/AgCl reference electrode with cross-aligned fiber architecture also shows a stable potential for over 3000 seconds, which can sustain 50% strain (**Supplementary Fig. 25**). Additionally, randomly oriented fibers showed lower initial conductivity (158 ± 6 S/cm) due to discontinuous conductive pathways and developed significant cracks when stretched (**Supplementary Fig. 26**), resulting in a further decrease in conductivity to 0.57 ± 0.03 S/cm at 100% strain (**Supplementary Fig. 27**). To evaluate the durability of E-cardiac, the platform was subjected to 100,000 cycles of 30% biaxial strain, equivalent to over 24 hours of continuous cardiac beating at a human physiological heart rate (**Supplementary Fig. 28**), revealing minimal resistance change (Δ*R* < 30 Ω). Post-cycling SEM characterization confirmed preservation of the cross-aligned fiber network architecture. (**Supplementary Fig. 29**). While minor localized fiber delamination (<5% of total fiber area) was observed at some high-stress junction regions, the overall microfiber structure remained intact with no significant fiber breakage, cracking, or network collapse, validating the robustness of the cross-aligned architecture.

The electrochemical performance of E-cardiac under deformation was first analyzed using K_3_[Fe(CN)_6_] solution (details given in **Supplementary Note 5**). Cyclic voltammetry at varying strain levels revealed nearly identical and symmetrical redox peaks, indicating excellent electrochemical stability during stretching. Quantitative analysis revealed almost constant peak current (variation less than 2.62%) and minimal variations in peak separation during stretching (**Supplementary Fig. 30**), while electrochemical impedance remained stable at 132 ± 6 Ω across 0-100% strain (**Supplementary Fig. 31**).

We further investigated H_2_O_2_ biosensing performance using amperometric methods at an optimized potential of -0.05 V, selected to balance sensitivity against baseline drift (**Supplementary Fig. 32**). E-cardiac demonstrates typical electrochemical biosensor characteristics, showing steady-state current increases proportional to H_2_O_2_ concentration, confirming its functionality as a reliable H_2_O_2_ sensor. Considering that sensing relies on the catalytic effect of encapsulated PB nanoparticles, we tuned the catalyst loading concentration to optimize the sensing performance: sensors with 2.5% PB nanoparticles achieved optimal sensitivity of 25.81 µA·cm^2^·mM^-1^, representing significant improvement compared to 2.35 µA·cm^2^·mM^-1^ for sensors without PB nanoparticles (**Supplementary Fig. 33**). At this optimized concentration, E-cardiac demonstrated a linear detection range from 0.5 µM to 300 µM with a limit of detection (LOD) of 0.38 µM, effectively covering the physiological range of H_2_O_2_ levels secreted by cells and enabling reliable monitoring in subsequent cellular and tissue experiments (**Figure 4c**). Moreover, E-cardiac exhibited high selectivity for H_2_O_2_ in the presence of common interferent metabolites found in tissue interstitial fluid, such as Na^+^, K^+^, Ca^2+^, Mg^2+^, NH ^+^ and potential redox-active species including H_2_S, NO, NADH, and NADPH (**Supplementary Fig. 34**).

Then, we evaluated the H_2_O_2_ sensing capabilities of E-cardiac under both static and dynamic deformation (details given in **Supplementary Note 6**). The sensor maintained consistent performance through both static conditions (0-50% strain, 10% increments) (**Supplementary Fig. 35**) and dynamic testing (0-50% strain at 0.02 mm/s) (upper panel in **Figure 4d**), exhibiting negligible response variation (≤15%) (left panel in **Figure 4e**). This stability is particularly significant given that arteries typically experience 5-10% strain under normal conditions and up to 20% under pathological conditions like hypertension^46^, while cardiac tissue undergoes ∼30% strain during beating^47^. In contrast, conventional PDMS/Au/PB electrodes exhibited significant signal degradation under both dynamic (lower panel in **Figure 4d**) and static stretching conditions (right panel in **Figure 4e**) at physiologically relevant strains, with nearly complete signal loss at a strain of 30%. The superior strain sustainability of E-cardiac is attributed to the nanoconfinement of catalysts within microarched fibers, preventing catalysts detachment during deformation while maintaining conductive pathways that resist strain effects. In contrast, the PDMS/Au/PB layer-by-layer configuration with PB directly exposed to solution suffers from both potential detachment of the active material and disruption of conductive pathways under deformation.

To elucidate the mechanical stability of E-cardiac electrochemical interfaces, we also developed theoretical framework correlating mechanical stability with the electrical properties underlying H_2_O_2_ sensing performance (**Supplementary Note 7**). The key insight is that biosensor performance under strain relies on the synergistic of electrical pathway integrity (maintaining low electrode resistance *R*_s_) and catalytic stability (sustaining robust electrochemical reactions at the tissue interface). Theoretical framework is illustrated in **Supplementary Figure 36**. Mechanical deformation of conventional multi-layered sensors leads to exponential resistance increase (*R*_s_ > 10³ Ω, *α* ≫ 1) and catalyst deactivation, drastically reducing performance (<10% sensitivity retention). In contrast, E-cardiac confines PB nanoparticles with gold nanoarches, where stress dissipates through particle rearrangement, preserving both conductivity (*α*=(*i*_0_·*R*_s,0_)/*η*_applied_=0.01) and catalyst stability (*k*_cat_(ε)≈*k*_cat_,_0_, *E*≈[*E*]_0_). The theorical framework predicts 98% sensitivity retention at 50% strain, consistent with experimental results. These quantitative agreements further explain why conventional biosensors fail on dynamic tissues while E-cardiac succeeds through synergistic combination of preserved electrical connectivity and biotransducer integrity.

Moreover, to show the versatility of this method, we further prepare different HRP-mimicking nanoparticles (CuO, MnO_2_, Fe_3_O_4_) encapsulated in E-cardiac and test their biosensing performance. TEM images show the successful encapsulation of various catalytic nanoparticles in fibers (**Supplementary Figure 37**). The obtained E-cardiac electrodes exhibited similar steady current increases with H_2_O_2_ addition (**Supplementary Fig. 38**), exhibiting biosensing sensitivities of 156.18 µA·cm^2^·mM^-1^, 53.56 µA·cm^2^·mM^-1^, 3.14 µA·cm^2^·mM^-1^ and LODs of 108.7 nM, 369.7 nM, and 4.96 µM, respectively. These results validate the versatility of the E-cardiac approach. To identify areas of myocardial ischemia during infarction and better understand the spatial distribution of oxidative stress during disease progression, we patterned the E-cardiac with a 2 × 4 array of working electrodes (each with a rectangular shape, with minimum achievable dimensions of 1.5 mm edge length) (**Supplementary Note 8**) with shared counter and reference electrodes. This design enables the capture of both the temporal and spatial distribution of H_2_O_2_ levels. Using a microfluidic system to mimic H_2_O_2_ flow, we demonstrated that each electrode recorded signals according to its relative position to the H_2_O_2_ source and flow direction, providing detailed spatiotemporal mapping of H_2_O_2_ distribution (**Figure 4f, g**).

We further investigated metabolic sensing capabilities by incorporating alternative sensing moieties into the hierarchical fiber architecture and evaluating their performance under different electrochemical mechanisms (**Supplementary Fig. 39**). PANI-encapsulated fibers exhibit a stable, potentiometric response over pH 4–9 (**Supplementary Fig. 39a, d**), enabling quantitative assessment of tissue acidosis. Glucose oxidase and lactate oxidase encapsulated fibers generate well-defined, concentration-dependent amperometric currents for lactate and glucose, respectively (**Supplementary Fig. 39b, c**), with strain-independent calibration curves showing <3% loss in sensitivity under 50% strain (**Supplementary Fig. 39e, f**). Collectively, these results demonstrate robust, deformation-tolerant multi-analyte sensing of pH, lactate, and glucose, establishing the platform as a generalizable interface for intraoperative metabolic monitoring.

### Oxidative stress profiling with E-cardiac across cells, tissues, and hearts

Before oxidative stress monitoring (with H_2_O_2_ as the indicator), we first assessed the biocompatibility and potential inflammatory responses of E-cardiac in endothelial cells and through dorsal subcutaneous implantation in rats. Comparable cell viability was obtained in the E-cardiac group relative to control groups for both 24 h CCK-8 assay (**Supplementary Fig. 40a**) and fluorescent imaging (**Supplementary Fig. 40b**), indicating good biocompatibility with mouse embryonic fibroblasts. Histological analysis at two weeks post-implantation showed minimal inflammation and tissue morphology comparable to that of the control group (**Supplementary Fig. 41**, **Supplementary Note 8**). Additionally, the E-cardiac demonstrated excellent gas permeability with water vapor transmission rate of ∼90 g/m^2^/h, attributed to their porous structure, which supports mass transfer during physiological processes (**Supplementary Fig. 42**).

Subsequently, we evaluated the capability of E-cardiac electrodes for H_2_O_2_ sensing across different biological scales, specifically at the cellular, tissue, and organ levels. Endothelial cells or cardiomyocytes generate H_2_O_2_ through oxidative metabolic pathways. In the vasculature, H_2_O_2_ plays a crucial role in regulating arteriolar tension, with high concentrations inducing vasodilation and low concentrations causing vasoconstriction. At the cardiac tissue level, oxidative stress is a key factor in the progression of myocardial dysfunction,^48^ particularly in cardiac diseases linked to hypertension^49^.

For *in vitro* H_2_O_2_ monitoring of cellular secretion, we used phorbol 12-myristate 13-acetate (PMA) as a stimulant to induce H_2_O_2_ generation from endothelial cells and cardiomyocytes, and monitored the H_2_O_2_ release using amperometry methods (**Supplementary Fig. 43a**). Upon PMA stimulation, both HUVECs and H9C2 exhibited a distinct increased reduction current (more negative current) within 10 seconds (**Supplementary Fig. 43b**), an effect not observed in control groups without PMA treatment. To further validate this observation, catalase, an enzyme that selectively decomposes H_2_O_2_, was added. This treatment resulted in comparable steady-state currents to those observed in the control group, further supporting the conclusion that the shifting in steady current is attributed to H_2_O_2_ release from the cells. These results suggest that the electrode effectively captures changes in extracellular hydrogen peroxide concentrations induced by PMA stimulation.

Next, we applied our E-cardiac to monitor H_2_O_2_ in isolated vascular ring segments of mouse aorta that primarily composed of endothelial cells^50^ (**Supplementary Fig. 44a**). E-cardiac successfully monitored the H_2_O_2_ release by mouse aorta, as indicated by an increased current upon PMA stimulation (**Supplementary Fig. 44b**). Catalase treatment effectively abolished the response current, confirming that the current signal originated from H_2_O_2_. These results validate E-cardiac’s capability to detect tissue-level oxidative stress, demonstrating its potential for monitoring vascular ROS dynamics in complex biological systems.

We further applied the E-cardiac to the wet beating heart surface throughout the cardiac motion, exhibiting seamless adhesion and continuous H_2_O_2_ monitoring that sustained to the cardiac motion (**Supplementary Video 3**). The electrochemical recording begins within seconds of attachment and continues throughout the surgical procedure. Notably, H_2_O_2_ sensing relies on the steady-state faradaic current, which is fundamentally distinct from the non-faradaic, periodic current fluctuations characteristic of ECG signals. We implemented a three-stage signal processing pipeline to extract steady-state current from raw signals, where dynamic Z-score thresholding removes high-amplitude ECG spikes, median filtering suppresses residual periodic interference, and Savitzky-Golay polynomial fitting isolates the H_2_O_2_-responsive faradaic current (**Supplementary Figure 45**). This orthogonal detection strategy enables real-time profiling of oxidative stress dynamics without interference from simultaneous cardiac electrical activity.

Subsequently, we investigated real-time H_2_O_2_ monitoring during IRI in rats (**Figure 5a-c**). IRI is an unavoidable complication of many cardiac surgeries, where coronary blood flow is intentionally interrupted (ischemia) and then restored (reperfusion), with each phase generating distinct oxidative stress patterns that are critical for surgical assessment. The ischemia model was established by ligating the left anterior descending artery (LAD) of a rat^51^ and followed by the reperfusion, after surgery, that release sutures to restore blood flow. Masson staining tissue sections revealed significant myocardial damage five days after the ligation (**Supplementary Fig. 45**). Representative amperometric traces (**Figure 5b**) illustrate distinct oxidative stress patterns across the three intraoperation scenario groups. The sham controls maintained a stable baseline current with minimal fluctuation (∼8.41 μM H_2_O_2_). The ischemia group exhibited a progressive current increase, indicating moderate H_2_O_2_ elevation (∼50.18 μM). In contrast, the ischemia-reperfusion group generated a substantially larger current surge upon restoration of blood flow, reflecting the characteristic oxidative burst with H_2_O_2_ levels reaching approximately 104.10 μM. To validate the sensing accuracy of E-cardiac, we employed the Amplex Red colorimetric assay as a gold-standard reference. H_2_O_2_ sensing results obtained from the E-cardiac sensor (red violin plots) demonstrated good correlation with the Amplex Red assay results (gray violin plots) across all experimental phases (n=8 for each group) (**Figure 5c**). This concordance confirms that the electrochemical platform accurately captures the physiological dynamics of oxidative stress throughout cardiac injury progression.

**Fig. 5.**
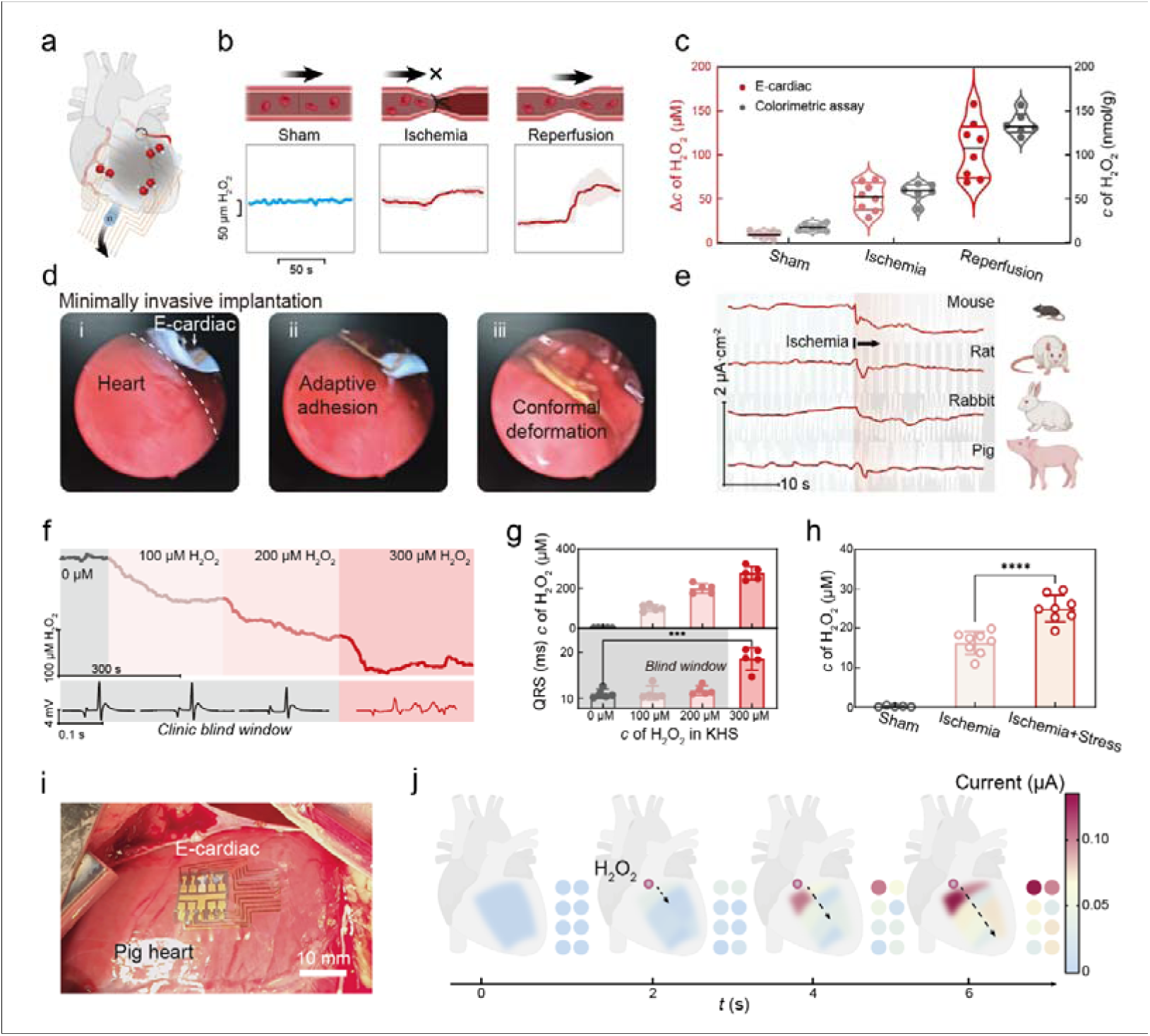
| E-cardiac validation in ischemia-reperfusion injury models: establishing clinical detection thresholds and high-fidelity monitoring on beating hearts. **a**, Schematic of E-cardiac for *in situ* H_2_O_2_ monitoring on the cardiac surface during surgery. **b**, Representative current-time curves detected by E-cardiac show progressive H_2_O_2_ elevation across ischemia-reperfusion injury phases. Ischemia generates higher ROS than sham baseline, while reperfusion induces maximal oxidative stress surge. **c**, Quantitative validation of E-cardiac H_2_O_2_ sensing, E-cardiac measurements (red violin plots) demonstrate excellent correlation with gold-standard colorimetric assays (gray violin plots), confirming progressive H_2_O_2_ elevation from sham to ischemia to reperfusion (n=8 hearts per group). **d**, Minimally invasive sensor deployment. The E-cardiac device is positioned on the beating heart using a catheter-based approach (i), where adaptive adhesion establishes immediate conformal contact (ii) and maintains stable integration throughout continuous cardiac cycles (iii). **e**, Cross-species validation: Real-time current responses from mouse, rat, rabbit, and pig hearts during ischemia demonstrate universal applicability across cardiac scales and beating frequencies. **f**, Langendorff-perfused heart model with graded H_2_O_2_ spiking (100, 200, 300 μM) simulate varying reperfusion injury severity. Simultaneous E-cardiac (red) and ECG (gray) recordings reveal that moderate oxidative stress (100-200 μM) maintains normal ECG despite ROS elevation (the “blind window”), while severe stress (300 μM) causes arrhythmia. **g**, Quantitative dose-response shows E-cardiac detects concentration-dependent H_2_O_2_ increases (top), including within the ECG blind window where QRS duration remains unchanged (bottom, 0-200 μM), and significant changes are observed at 300 μM (p < 0.001), establishing early detection advantage before functional compromise. **h**, Mechanical interference assessment: Significantly lower H_2_O_2_ detected in ischemic hearts with external stress versus unstressed controls (unpaired t-test, ****p < 0.0001, n=8), confirming rigid sensors induce mechanical artifacts. **i**, **j,** Image of the 2 × 4 E-cardiac sensor array on the dynamic heart (i) and time-resolved imaging (j) of H_2_O_2_ added to simulate IRI.

To demonstrate the clinical translational potential for minimally invasive cardiac surgery, we investigate the deployment of E-cardiac through small incisions in both rat (<1 cm) (**Figure 5d, Supplementary Fig. 47, Supplementary Video 4**) and pig (<3 cm) models (**Supplementary Fig. 48, Supplementary Video 5**). The ultrathin E-cardiac sensor (∼460 nm) was positioned on the beating heart (i), achieved adaptive adhesion to the epicardial surface within seconds (ii), and maintained conformal deformation throughout cardiac cycles (iii). These characteristics expand applications in video-assisted thoracoscopic surgery (VATS), robot-assisted procedures, and pericardial-access interventions.

After that, we confirmed the feasibility in multiple animal models including mice, rabbit and pig hearts using our E-cardiac. The critical role of PB nanoparticles in H_2_O_2_ sensing was confirmed through control experiments: E-cardiac without PB nanoparticles showed negligible current response during ischemia (**Supplementary Fig. 49**). Similar results were reproduced in rat, rabbit and pig hearts (**Figure 5e)**, where ischemia induced significant current changes with estimated H_2_O_2_ burst concentrations reaching tens of micromolar, while no response was observed in sham groups (**Figure 5a**).

The Langendorff isolated hearts were further utilized to simulate graded perfusion states to mimic varying IRI severity independent of systemic confounders. This approach circumvents inter-animal variability inherent to partial coronary occlusion while simulating intrinsic, physiologically relevant oxidative stress thresholds. E-cardiac sensors were attached to perfused rat hearts maintained at 37°C with oxygenated buffer (**Supplementary Fig. 50**). We precisely modulated perfusate H_2_O_2_ concentrations (100, 200, 300 μM) to recapitulate oxidative stress gradients occurring during cardiac procedures with variable perfusion adequacy, while concurrently acquiring real-time amperometric measurements. Representative current-time profiles establish a stable baseline during blank perfusion. Sequential 100 μM H_2_O_2_ injections yield highly reproducible, stepwise current escalations (**Figure 5f**), validating the specificity for H_2_O_2_ sensing.

Quantification across multiple hearts (n=5) shows linear, dose-dependent response with accurate concentration measurements (r² > 0.98). Critically, we simultaneously recorded ECG throughout H_2_O_2_ exposure to correlate metabolic and electrophysiological responses. The E-cardiac platform substantially extends the clinically actionable window for detecting suboptimal yet surgically correctable perfusion, resolving moderate elevations in H_2_O_2_ (100–200 μM) under conditions where conventional electrophysiologic indices—heart rate, PR interval, and QRS duration—remain unchanged (p > 0.05, **Figure 5g**). Elevated H_2_O_2_ levels (≥300 μM) cause severe dysfunction: prolonged PR interval and QRS duration (p<0.001), arrhythmia, and cardiac arrest risk, indicating transition to potentially irreversible injury. Synchronized ECG comparisons confirm that the E-cardiac detects escalating oxidative stress, providing an early metabolic warning window well before alterations in cardiac electrophysiology occur. This early metabolic warning identifies reversible injury, enabling preemptive surgical intervention before functional compromise occurs.

The mechanical invisibility of E-cardiac was validated in Langendorff perfused hearts by measuring left ventricular systolic blood pressure (LVSbp) and pulse pressure, indicators of contractile performance (**Supplementary Figure 51**). E-cardiac attachment produced negligible hemodynamic impact, yielding LVSbp and pulse pressure values statistically equivalent to controls (LVSbp: 133.3 ± 1.5 vs. 131.8 ± 2.3 mmHg, pulse pressure: 54.2 ± 1.3 vs. 52.7 ± 1.3 mmHg, p>0.05). Conversely, conventional PDMS film caused significant mechanical restriction, reducing both parameters (LVSbp: 124.0 ± 1.2 mmHg, pulse pressure: 46.3 ± 1.3 mmHg, p<0.01). To quantitatively validate the impact of mechanical stress on ROS production, we applied external pressure (50 kPa, pressure estimated by our simulation for achieving a good contact between biosensors and cardiac tissues) to the mouse heart (**Figure 5h**). An obvious increase in H_2_O_2_ production than only ischemia was observed, further demonstrating the advantage of our stress-deconcentrated strategy that could minimize the ROS interferences that induced by mechanical stress. These results confirm the mechanical adaptation of E-cardiac that preserves native tissue function, avoiding the mechanical constraint imposed by stiffer materials.

To facilitate clinical translation of E-cardiac, we evaluated it’s performance including intraoperative stability and straightforward removal. *In vivo* stability was assessed through 1-hour monitoring on beating rat hearts (∼24,000 cycles at 400 bpm), maintaining structural integrity with no delamination or cracking (**Supplementary Video 6**). This corresponds to ∼6.7 hours at human heart rate (60 bpm), exceeding typical surgical durations (2-5 hours). Despite strong adhesion for stable monitoring, E-cardiac is easily removed within seconds via gentle wiping with moistened cotton swab (**Supplementary Figure 52 and Supplementary Video 7**). E-cardiac removal is atraumatic, with no tissue damage, residual material, or epicardial disruption, allowing repositioning if needed and complete extraction post-procedure. This temporary placement-and-removal workflow suits acute intraoperative applications, functioning as transient electronics for surgical guidance.

Spatial mapping of ROS distribution during myocardial events could provide valuable insights for targeted therapeutic interventions. We further applied the 2 × 4 E-cardiac array (**Figure 5i**) for mapping H_2_O_2_ levels on a rat heart with exogenous H_2_O_2_ mimicking oxidative stress during cardiac ischemia (**Figure 5j**). The E-cardiac array facilitated the real-time visualization of H_2_O_2_ distributions across cardiac regions, highlighting its capacity for high spatial resolution. Moreover, the modular design of E-cardiac enables scalable assembly into high density configurations (**Supplementary Figure 53**). A 60-sensor high-density array was deployed on pig heart demonstrated enhanced spatiotemporal resolution for precise ischemic zone localization. Crucially, the E-cardiac enables ROS mapping without inducing mechanical disruption to cardiac function, making it a valuable tool for monitoring disease progression and assessing the efficacy of targeted therapeutic interventions within the infarcted myocardium.

### Conclusion

In this work, we present the E-cardiac platform, featured by the hierarchical mechanical-adaptation architecture that enables mechanical-artifact-free catalytic electrochemical sensing on highly deformable, continuously beating heart. This capability could address a critical and previously unmet clinical need for real-time oxidative stress monitoring during cardiac surgery, where ischemia-reperfusion injury generates ROS before conventional biomarkers or ECG abnormalities appear. The interfacial stress deconcentration operates through macro-scale biofluid-mediated conformable contact, micro-scale fiber reorganization, and nanoscale enzymatic confinement. Simulation data demonstrates interfacial stress below the kPa threshold, and cellular experiments confirm absence of mechanosensitive PIEZO channel activation, validating the mechanically invisible design. E-cardiac achieves wide linear H_2_O_2_ detection (0.5 μM to 300 μM) with low detection limit (0.38 μM), robust biosensing performance under 50% strain, high endurability with prolonged mechanical cycling (100,000 cycles), and transient biointegration—the device can be safely removed or naturally degraded post-surgery. Deployment versatility was demonstrated through both open-heart and minimally invasive approaches in rat and pig models. Validated across cardiomyocytes, tissues, *in vivo* ischemia-reperfusion models, E-cardiac demonstrated reliable oxidative stress detection with excellent correlation to gold-standard Amplex Red colorimetric assay, confirming quantitative accuracy across physiological to pathological H_2_O_2_ ranges. In *ex vivo* Langendorff-perfused rat hearts with simultaneous ECG monitoring, E-cardiac successfully detected escalating oxidative stress within the “ECG blind window” (100-200 μM H_2_O_2_) where cardiac electrical function remained normal, providing early intervention opportunities before irreversible tissue damage. By addressing critical mechanotransduction-related artifacts, this advancement enables molecular-level detection directly on the beating heart, offering significant potential for real-time revelation of disease propagation mechanisms at the molecular level *in vivo*.

## Methods

### Materials and reagents

Chitosan, K_3_[Fe(CN)_6_], ethanol, Polyvinylpyrrolidone, PVA1788, H_2_O_2_, and catalase were obtained from Aladdin. HCl was purchased from Sinorengent. Silver/silver chloride (60/40) paste was obtained from Sigma-Aldrich. PDMS was sourced from Dow Corning, and Ecoflex 00-30 was purchased from Smooth-On Inc. Cell culture reagents, including PBS, DMEM, and FBS, were obtained from Keygentec. Calcein AM, DAPI, and DCFH-DA were purchased from Beyotime, while Fluo-4 AM was sourced from Yeasen. Surgical consumables, such as surgical instruments and sutures, were purchased from RWD. All reagents were used without further purification.

### Fabrication of E-cardiac

Cross-aligned fiber networks were fabricated with radial electrospinning methods. Briefly, pre-synthesized mimic enzymes (PB nanoparticles; synthesis details in Supporting Information) were dispersed in an electrospinning precursor solution consisting of 10 wt% PVA and 1 wt% chitosan in deionized water. The electrospinning was conducted using a Biofabrication apparatus with a 22 G metal needle, collecting fibers on a high-speed rotating drum (2800 rpm). The electrospinning parameters were optimized at 15 kV applied voltage (12 kV positive, - 3 kV negative), 15 cm working distance, and 5 μ min^−1^ flow rate for 30 minutes. The same encapsulation strategy was applied to other biotransducers, including GOx, LOx, PANI, and alternative catalytic nanoparticles, by substituting the corresponding functional component in the precursor solution. A 50 nm-thick Al_2_O_3_ layer was first deposited on the wire regions to provide isolation, followed by a patterned 70 nm-thick gold layer through thermal evaporation using a shadow mask. Finally, an additional Al_2_O_3_ layer was deposited for complete insulation.

### Contact stress analysis through mechanical modeling

The heart surface is analogously modeled after the skin and is represented by a sinusoidal profile: *y*_0_ (*x*) = *h*_0_[1+ cos(2*π x* / *λ*_0_)], where *h*_0_ and *λ*_0_ are the semiamplitude and wavelength of the undeformed heart surface, respectively. Considering the total energy of the E-cardiac–heart system, which includes contributions from E-cardiac bending, heart elasticity, and interfacial adhesion, the critical wavelength *λ*_0_ that enables a thin membrane to conformally adhere is given by:

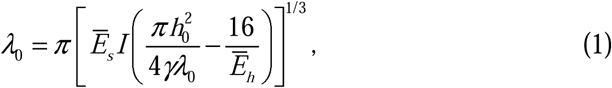

where *γ* is the work of adhesion, *E̅_s_* and *E̅_h_* are the plain-strain modulus of the E-cardiac and heart, respectively.

The maximum contact stress during the dynamic deformation of the heart caused by beating is given by:

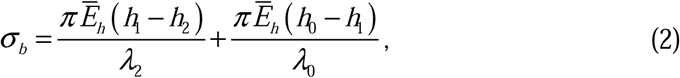

where *h*_1_ = *h*_0_*ξ*, *h*_2_ = *h*_1_ (1− *ε_b_*), *λ*_2_ = *λ*_0_ (1+ *ε_b_*). Here, *ε_b_* represents the equivalent dynamic strain of the heart, and *ξ* is a normalized amplitude factor obtained by solving the following equation:

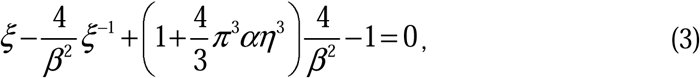

where *β*=2*πh*_0_ / *λ*_0_, *α* = *E̅_s_* / *E̅_h_* and *η* =*t* / *λ*_0_. Here, *t* is the thickness of the membrane. Details of the theoretical analysis are provided in the Supporting Information.

### ROS detection in HUVECs under mechanical stress

HUVECs were cultured in Endothelial Cell Medium supplemented with fetal bovine serum, Endothelial Cell Growth Supplement (ECGS, ScienCell Cat. #1052), and penicillin/streptomycin in a humidified incubator (95% air, 5% CO_2_). For ROS detection, cells were cultured in six-well plates and either interfaced with E-cardiac probes or subjected to mechanical stress (5.0 kPa, applied via PDMS-coated weight). Intracellular ROS levels were assessed using DCFH-DA (10 μM, 25-minute incubation) after 2-hour treatment. ROS levels were analyzed using fluorescence microscopy (Olympus) and flow cytometry (Beckman Coulter).

### Detection of cell- and tissue-derived H_2_O_2_

HUVECs were seeded on E-cardiac probes at ∼5 × 10^6^ cells/mL for cellular H_2_O_2_ detection. H9C2 cardiomyocytes were also tested under similar conditions. For aortic ring assays, thoracic aortas were harvested from isoflurane-anesthetized mice following standard aseptic procedures. After cleaning surrounding tissues, aortas were sectioned into 1-mm rings and placed on E-cardiac probes.

H_2_O_2_ generation was stimulated using PMA (1 μg/mL). Catalase (100 U/mL) was used as a negative control to confirm H_2_O_2_-specific responses by enzymatically depleting H_2_O_2_. Electrochemical measurements were performed using standard connections to the electrochemical workstation.

### Method of induction of IRI and ROS sensing

Myocardial ischemia-reperfusion injury (IRI) was established in Sprague-Dawley (SD) rats utilizing a reversible left anterior descending (LAD) coronary artery ligation model. Animals were anesthetized, and the heart was accessed via a left thoracotomy. The LAD was ligated using a 6-0 silk suture secured with a slipknot, with successful occlusion visually confirmed by the immediate pallor of the anterior left ventricular wall. Following a 15-min ischemic period, the slipknot was released to initiate myocardial reperfusion.

To demonstrate the cross-species scalability of the E-cardiac platform under ischemic conditions, isolated myocardial ischemia was subsequently induced in murine (C57BL/6), rabbit, and porcine models. LAD ligation was executed using analogous surgical principles scaled to respective anatomical specifications. Standardized post-operative protocols, including analgesia and physiological monitoring, were administered across all animals.

For ROS monitoring, the E-cardiac biosensor was placed on the cardiac surface, where it spontaneously adhered through physiological fluid-driven interfacial contact. E-cardiac Signal readout is achieved via a PET/Au array interconnect, which bridges the biosensor to measurement electronics through an anisotropic conductive film. To evaluate sensing performance under mechanical interference, controlled pressure (up to 50 kPa) was applied during ROS monitoring using a force transducer system. All animal procedures were approved by the Institutional Animal Care and Use Committee of Nanjing Medical University (IACUC2404016), and Nanjing University (2024AE05007).

### Langendorff models with H_2_O_2_ spiked testing under heart beating

Following anesthesia, rats received an intraperitoneal injection of sodium heparin solution to achieve systemic anticoagulation. Animals were subsequently euthanized by cervical dislocation. Each rat was placed in the supine position on a surgical platform, and the thoracic cavity was rapidly opened. The heart was excised intact and immediately immersed in ice-cold (4°C) Krebs-Henseleit (KH) buffer. After trimming the aortic root, the aorta was cannulated onto the aortic cannula of a Langendorff perfusion system and subjected to retrograde perfusion with KH solution maintained at 37°C and continuously gassed with 95% O_2_/5% CO_2_, until spontaneous rhythmic contractions were restored. The heart was then allowed to stabilize for 15 minutes.

Subsequently, ECG recording electrodes were positioned on the epicardial surface of the left ventricular wall. In parallel, the E-cardiac biosensor was placed onto the epicardial surface of the left ventricular wall. Once stable baseline signals were established, hydrogen peroxide (H_2_O_2_) solutions of incrementally increasing final concentrations were introduced into the perfusate reservoir. ECG signals and the electrochemical response of the E-cardiac biosensor were recorded simultaneously and continuously throughout the experiment.

All animal procedures were approved by the Institutional Animal Care and Use Committee of Henan Key Laboratory of Cardiac Electrophysiology (LLSC0025)

## Acknowledgements

We thank the financial support from Natural Science Foundation for Excellent Young Scholars (62322108), the National Natural Science Foundation of China (62288102, 62235008), Natural Science Foundation for Young Scholars (62201286, 62301283, 22405131), the Program of Jiangsu Specially-Appointed Professor, Science Foundation of Nanjing University of Post and Telecommunications (NY221004, NY222099), Basic Research Program of Jiangsu (BK20253006, BK20243057), and the Agency for Science, Technology and Research (A*STAR) under its MTC Programmatic Funding Scheme (M23L8b0049) Scent Digitalization and Computation (SDC) Programme.

## Author contributions

B.-W. Y., T. W., X.D. C. and L.H. W. designed research; B.-W. Y., J.- W. W., Z.C., L.- X. X., L. X., and H. Z. performed research; D. W., Y.-X. W., C. C. and Q.-L. L. conducted theory and simulation, B.-W. Y., Y.-Y. X., X.-J. H., P.-Q. C. and B.-H. H. contributed animal models; B.-W. Y., X. Z. and T. L. contributed new reagents/analytic tools; B.-W. Y., N. C. and T. W. analyzed data; and B.-W. Y., T. W., X.D. C. and L.-H. W. wrote/revise the paper. X.G. and H. L. contributed to the minimal invasive implants to pig; Y. D., G. H, G. W. and Z. Z. contribute to the Langendorff models.

## Additional information

Supplementary information is available in the online version of the paper. Reprints and permissions information is available online at www.nature.com/reprints. Correspondence and requests for materials should be addressed to L.-H. W.

